# A knock-in *Six2Cre* line reveals transient interstitial potential in nephron progenitors

**DOI:** 10.64898/2026.02.04.703893

**Authors:** Azadeh Haghighitalab, Fariba Nosrati, Zeinab Dehghani-Ghobadi, Mohammed Sayed, Christopher Ahn, Yueh-Chiang Hu, Eunah Chung, Hee-Woong Lim, Joo-Seop Park

## Abstract

The developmental relationship between nephron progenitors and the renal interstitium remains unresolved, in part due to limitations of existing lineage tracing tools. The widely used transgenic *Six2TGC* line, which is routinely employed to target the nephron lineage, exhibits mosaic recombination and altered progenitor dynamics. To overcome these shortcomings, we generate a knock-in *Six2Cre* mouse allele that faithfully recapitulates endogenous *Six2* expression, preserves nephron endowment, and achieves near-complete, non-mosaic recombination. Side-by-side lineage tracing with *Six2Cre* and *Six2TGC*, combined with RNA velocity analysis of single-cell RNA-sequencing datasets, reveals a brief interval around embryonic day 11 during which *Six2*-expressing mesenchymal nephron progenitors contribute to the renal interstitium. This contribution is transient and stage-restricted. These findings reveal an early dual potential within nephron progenitors and define a precise developmental window for dissecting mechanisms that coordinate nephron-interstitium integration.

## Introduction

The mammalian kidney develops through the coordinated interaction of multiple progenitor populations that give rise to the nephron, collecting duct, vasculature, and interstitium^1, 2^. Among these, mesenchymal nephron progenitor cells (mNPs) are essential for nephrogenesis, as they generate all epithelial components of the nephron, including podocytes and tubular segments^3^. Proper regulation of mNP self-renewal and differentiation is required to sustain nephron formation throughout development and to establish final nephron endowment^4, 5^.

The transcription factor Six2 is a defining marker of mNPs and plays a central role in maintaining their progenitor state. Loss of Six2 leads to premature differentiation and depletion of the nephron progenitor pool, suggesting that *Six2* expression is required for preserving nephrogenic potential^4, 6^. Because of its restricted expression pattern and functional importance, Six2 has been widely used as a genetic entry point to study nephron progenitor behavior, lineage relationships, and gene function during kidney development^3, 7^.

In contrast to mNPs, the renal interstitium is thought to arise from a distinct population of stromal, or interstitial, progenitors that surround the nephron progenitor niche and the collecting duct^8, 9, 10^. These progenitors give rise to multiple interstitial cell types that play essential roles in kidney patterning, vascular development, and nephron differentiation. During early kidney development, these populations are molecularly defined by transcription factors such as Foxd1^8^, which marks cortical stromal progenitors, and Tbx18^9, 10^, which marks ureteric interstitial progenitors. Genetic lineage tracing studies have therefore suggested a model in which nephron progenitors and stromal progenitors represent separate and lineage-restricted compartments^3, 8^.

Consistent with this model, *Six2* expression has been widely interpreted as a marker of commitment to the nephron lineage, with no contribution to interstitial cell fates^3^. Genetic lineage tracing using *Six2*-driven Cre recombinase has been instrumental in defining nephron progenitor contributions^3^. However, the most commonly used bacterial artificial chromosome (BAC) transgenic *Six2* Cre line, *Six2TGC*, exhibits several limitations, including mosaic recombination and reduced nephron endowment^3, 7, 11, 12, 13, 14^. Such defects complicate interpretation of lineage tracing experiments and can confound functional studies that rely on efficient and faithful recombination within the nephron lineage.

To overcome these limitations, we generated a novel *Six2* knock-in Cre mouse allele that preserves endogenous *Six2* regulation and enables robust, non-mosaic recombination. Using this allele for lineage tracing, we unexpectedly found that *Six2*-expressing mNPs contribute to the renal interstitium during early kidney development. This finding is supported by complementary lineage tracing using *Six2TGC* and by RNA velocity analysis of published singlel-cell RNAl-seq datasets. Together, these results reveal an unanticipated developmental relationship between nephron and interstitial lineages, challenging the prevailing view that *Six2* expression marks exclusive commitment to the nephron lineage.

## Results

### KnockLin *Six2Cre* reproduces endogenous *Six2* expression

We generated a *Six2* knock-in Cre (*Six2Cre*) mouse line using CRISPR/Cas9-mediated genome editing. A validated single guide RNA (sgRNA) targeting the *Six2* locus was used to induce double-stranded breaks in fertilized C57BL/6J eggs, which were then injected with Cas9 protein, sgRNA, and a donor plasmid containing a *T2A-NLS-GFP-Cre* cassette^15^, where the NLS (nuclear localization signal) directs the GFP-Cre fusion protein to the nucleus, flanked by homology arms (Supplementary Fig. 1). Following embryo transfer, pups were screened by long-rage PCR and Sanger sequencing, confirming the correct integration of the cassette immediately downstream of the *Six2* stop codon. Because insertion of the *T2A-NLS-GFP-Cre* cassette preserves the *Six2* coding sequence^15^, the *Six2Cre* allele can be maintained in the homozygous state, making mouse breeding and genetic crosses highly efficient.

We examined *GFP-Cre* expression in wild-type, *Six2Cre*, and *Six2TGC* mouse embryos and kidneys (Fig. 1). *Six2* has been reported to be expressed in the head mesoderm as well as in the tendons and ligaments of the developing limb^16, 17^. At embryonic day 13.5 (E13.5), *Six2Cre* embryos faithfully recapitulated this endogenous expression pattern, with GFP-Cre detected in the same anatomical domains (Fig. 1a). In contrast, *Six2TGC* embryos exhibited reduced GFP-Cre signal in the head mesoderm and lacked detectable GFP-Cre in the developing limb. In newborn (P0) kidneys, both *Six2TGC* and *Six2Cre* displayed the characteristic horseshoe-shaped GFP signal in the nephrogenic niches corresponding to mNPs, although the GFP signal was stronger in *Six2TGC* kidneys (Fig. 1b,c), likely reflecting the presence of multiple copies of the *GFP-Cre* transgene^13^. Notably, *Six2Cre* kidney did not show the size reduction observed in *Six2TGC* kidneys (Fig. 1b)^7, 13, 18, 19^.

**Fig. 1:**
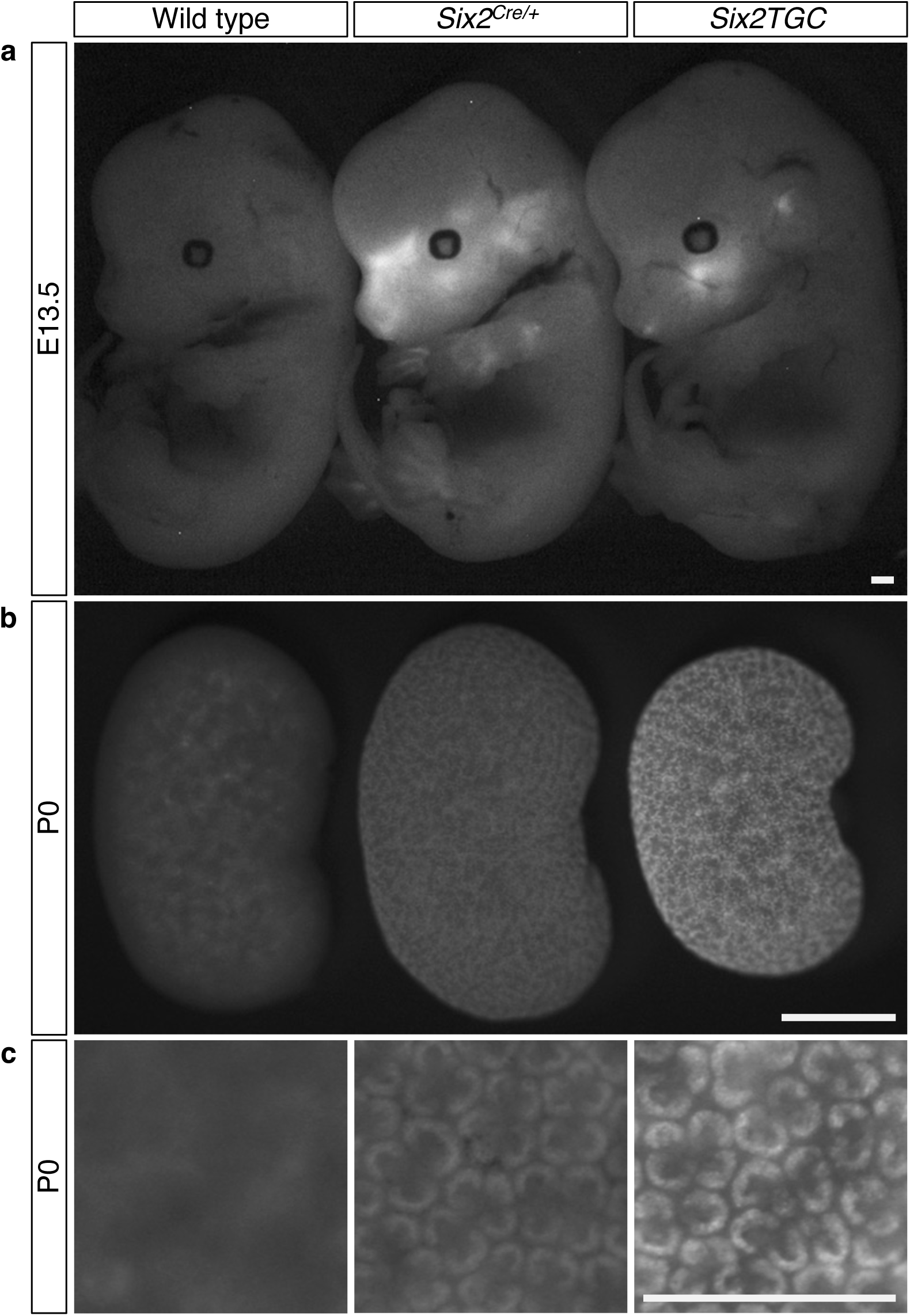
***GFP-Cre* expression in *Six2Cre* and *Six2TGC* mouse embryos and kidneys** (**a**) Both the knock-in *Six2Cre* and BAC transgenic *Six2TGC* alleles drive express of a GFP-Cre fusion protein. In *Six2Cre* embryos, *GFP-Cre* expression faithfully recapitulates endogenous *Six2* expression in the head mesoderm and in tendons and ligaments of the developing limb. In contrast, *Six2TGC* embryos show reduced GFP signal in the head mesoderm and no detectable expression in limbs. Stage: E13.5. (**b**) *Six2TGC*-positive kidneys are smaller than *Six2Cre*-positive and wild type kidneys. Stage: P0. (**c**) Both *Six2Cre*- and *Six2TGC*-positive kidneys show *GFP-Cre* expression outlining the horseshoe-shaped nephrogenic niches corresponding to mesenchymal nephron progenitors, with *Six2TGC* kidneys displaying higher GFP signal intensity. Stage: P0. (**a**-**c**) Scale bar, 0.5 mm.

### Transcriptional comparison of *Six2TGC* and *Six2Cre* mNPs

Despite lower GFP signal intensity in *Six2Cre* kidneys compared with *Six2TGC* (Fig. 1b,c), welll-defined GFPl-positive populations were readily isolated by fluorescencel-activated cell sorting (FACS) (Supplementary Fig. 2a), enabling downstream transcriptional analyses.

To assess how expression of *Six2TGC* and *Six2Cre* influences the transcriptional state of mNPs, we performed bulk RNA sequencing on FACS-isolated mNPs from *Six2TGC* and *Six2Cre* kidneys. Consistent with previous reports^13^, *Six2TGC*-expressing mNPs showed ectopic expression of *Six3*, a known artifact of the BAC transgenic line (Supplementary Fig. 2b). In contrast, *Six2Cre*-expressing mNPs showed no detectable *Six3* expression, indicating that the knock-in allele more faithfully preserves the endogenous transcriptional program.

We next examined the expression of key transcriptional regulators required for mNP identity and maintenance. Canonical mNP genes, including *Eya1*, *Osr1*, *Pax2*, *Wt1*, and *Sall1*, showed no significant differences in expression between *Six2TGC*- and *Six2Cre*-expressing mNPs. However, *Cited1*, a more stringent marker of self-renewing mNPs^20, 21, 22^, was expressed at significantly higher levels in *Six2Cre*-expressing mNPs than in *Six2TGC*-expressing mNPs (Supplementary Fig. 2b). Consistent with previous findings that *Cited1* expression is regulated by FGF signaling^23^, *Cited1* expression positively correlated with *Fgf8* expression in mNPs.

Although *Fgf8* expression is classically associated with mNP differentiation^24, 25^, analysis of previously published scRNA-seq datasets (GSE178263)^26^ from developing mouse kidneys indicates that a subset of mNPs express *Fgf8* (Supplementary Fig. 2c). In contrast, genes associated with mesenchymal-to-epithelial transition, including *Wnt4*, *Jag1*, *Notch1*, and *Lhx1*, were expressed at lower levels in *Six2Cre*-expressing mNPs than in *Six2TGC*-expressing mNPs (Supplementary Fig. 2b). Together, these transcriptional features suggest that *Six2Cre* more effectively preserves the progenitor state of mNPs compared with *Six2TGC*.

### *Six2Cre* enables efficient nephron gene deletion

We assessed recombination efficiency of *Six2TGC* and *Six2Cre* using proximal tubule cells as a proxy, as their origin from mNPs and numerical abundance provide a sensitive readout of Cre activity in the nephron progenitor lineage. Specifically, we examined *Six2TGC*-mediated activation of the *Rosa26-Sun1* reporter, which upon recombination produces a GFP fusion protein localized to the nuclear membrane^27^, and *Six2Cre*-mediated deletion of *Hnf4a*, a transcription factor required for the formation of mature proximal tubule cells^14, 28^.

In newborn kidneys (P0), lineage tracing using *Six2TGC* revealed incomplete targeting of proximal tubule cells, as indicated by the presence of Hnf4a+ cells that lacked reporter activation (Fig. 2a). Consistent with this mosaic recombination, conditional deletion of *Hnf4a* using *Six2TGC* has been shown to allow the persistence of Hnf4a+ proximal tubule cells^14^.

**Fig. 2:**
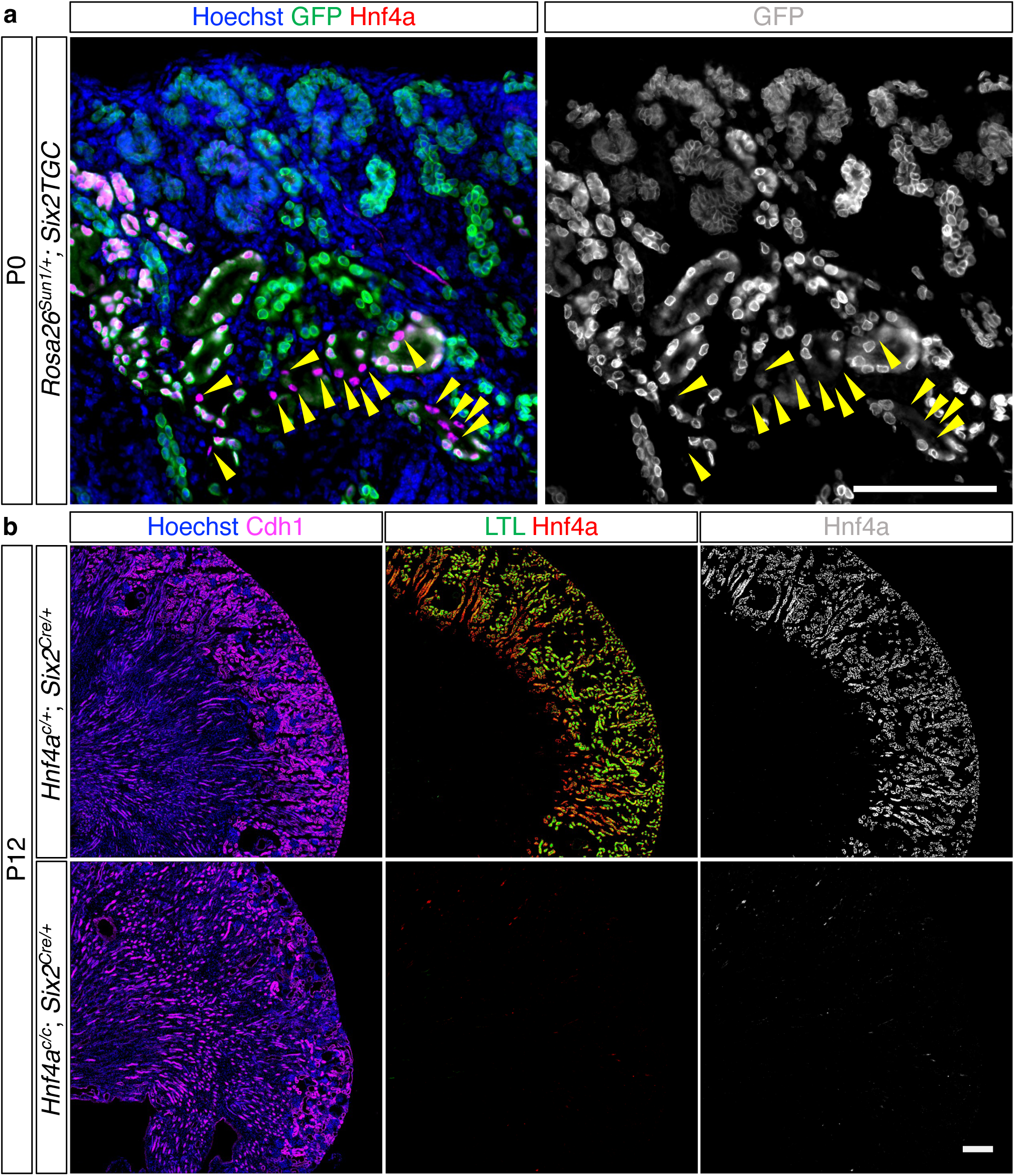
**Differential targeting efficiency of proximal tubule cells by *Six2TGC* and *Six2Cre*** (**a**) Lineage tracing reveals incompletely targeting of Hnf4a+ proximal tubule cells by *Six2TGC*, with Rosa26-Sun1 reporter-negative cells indicated by yellow arrowheads. Stage: P0. (**b**) In contrast, *Six2Cre*-mediated deletion of *Hnf4a* results in complete loss of Hnf4a+ proximal tubule cells. LTL marks mature proximal tubules, and Cdh1 labels epithelial cells. Stage: P12. (**a, b**) Scale bar, 200 µm.

In contrast, *Six2Cre*-mediated recombination resulted in uniform targeting of the nephron lineage. Deletion of *Hnf4a* using *Six2Cre* led to a near-complete loss of Hnf4a+ cells in the kidney (Fig. 2b). Consistent with the essential role of Hnf4a in proximal tubule maturation^14, 28^, loss of Hnf4a was accompanied by a complete loss of Lotus tetragonolobus lectin (LTL) staining, which marks mature proximal tubules, whereas the epithelial marker Cdh1 was preserved. These findings demonstrate that *Six2Cre* enables efficient and complete recombination throughout the nephron lineage, overcoming the mosaic targeting observed with *Six2TGC*.

#### *Six2Cre* also targets subsets of interstitial cells

While characterizing the lineage specificity of *Six2Cre* using the *Rosa26-Sun1* reporter, we unexpectedly observed GFP-positive cells outside the epithelial nephron lineage. This unanticipated observation prompted us to examine whether *Six2Cre* also targets interstitial cell populations. To address this possibility, we analyzed kidneys using Rosa26-Sun1-based lineage tracing in combination with markers of nephron progenitors, epithelial cells, and interstitial cells.

Immunostaining for Six2 and cytokeratin confirmed that *Six2* expression was restricted to mNPs, whereas cytokeratin marked collecting duct epithelial cells (Fig. 3a). Lineage tracing showed that *Six2Cre* efficiently labeled all Six2+ cells (Fig. 3a), consistent with robust targeting of the nephron lineage (Fig. 2b).

**Fig. 3:**
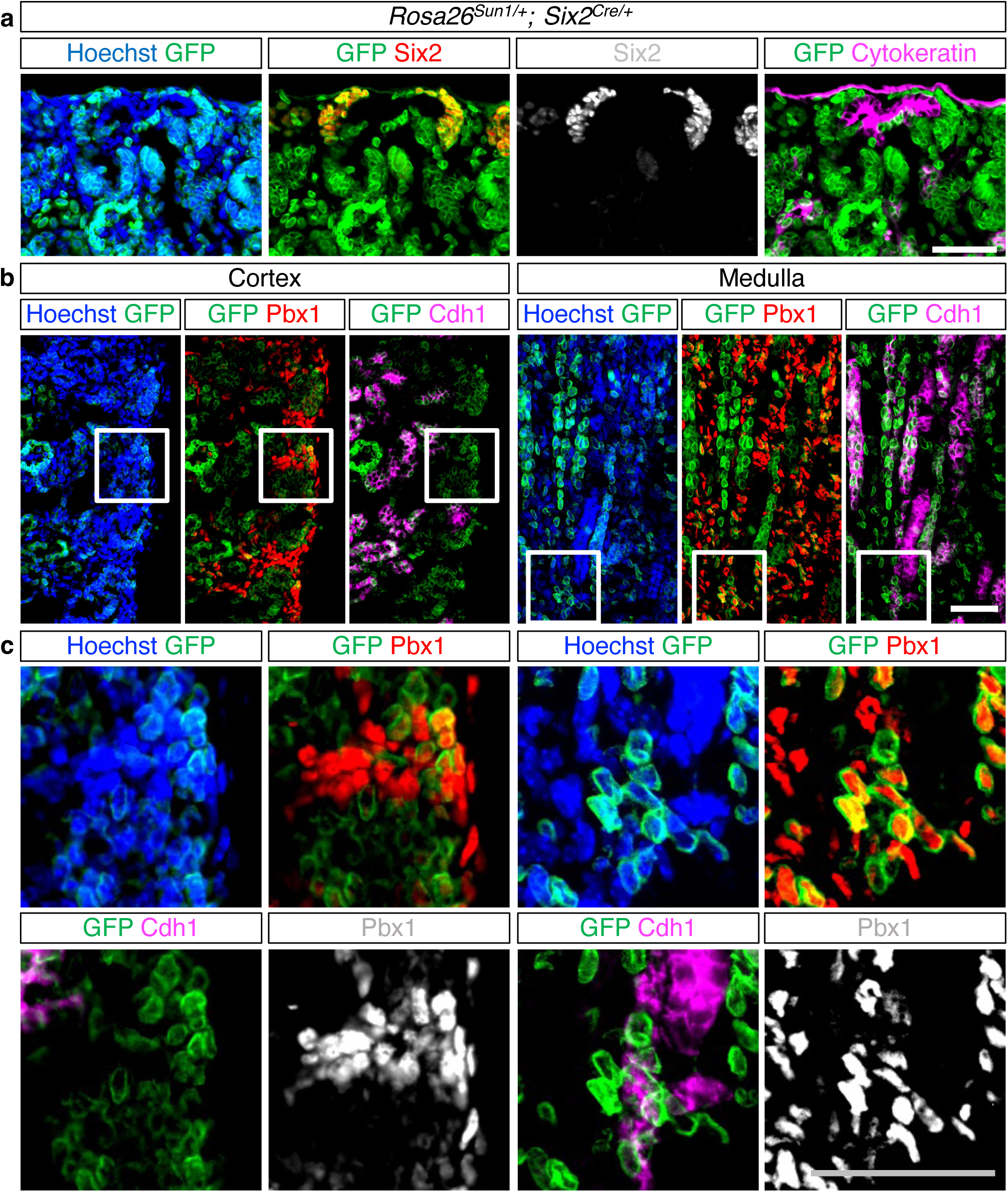
***Six2Cre* targets a subset of interstitial cells in addition to the nephron lineage** (**a**) Six2 marks mesenchymal nephron progenitors, whereas cytokeratin labels the collecting duct epithelium. *Six2Cre* lineage tracing labels all Six2+ nephron progenitors. (**b**) Cdh1 labels epithelial cells, while *Pbx1* is highly expressed in interstitial cells and weakly expressed in mesenchymal nephron progenitors. A subset of Pbx1+ Cdh1- interstitial cells is labeled by *Six2Cre*-activated Rosa26-Sun1 reporter in both cortical and medullary regions. White boxes indicate regions shown at higher magnification in (**c**). (**a-c**) Stage: P1; Scale bar, 50 µm.

Next, we examined *Rosa26-Sun1* reporter activation relative to Cdh1 and Pbx1. Cdh1 labels renal epithelial cells, whereas Pbx1 marks interstitial cells^29, 30^. Accordingly, interstitial cells were defined by strong Pbx1 signal in the absence of Cdh1 signal. Using these criteria, we identified GFP-positive cells within both cortical and medullary interstitial compartments (Fig. 3b,c). Notably, GFP labeling was confined to a subset of interstitial cells, indicating that *Six2Cre* does not target the entire interstitium (Supplementary Figure 3).

### Early Six2+ mNPs generate nephron and interstitial cells

To determine when *Six2*-expressing cells contribute to the interstitial lineage, we performed lineage tracing at earlier stages of kidney development using the *Rosa26-Sun1* reporter activated by *Six2Cre*. At embryonic day 10 (E10), all Six2+ Pax2+ mNPs cells were uniformly labeled by GFP and were positioned adjacent to the nephric duct epithelium, which exhibited strong Pax2 signal (Fig. 4a). At this stage, GFP labeling was largely confined to Six2+ mNPs, indicating that *Six2Cre* efficiently marks the early nephron progenitor population.

**Fig. 4:**
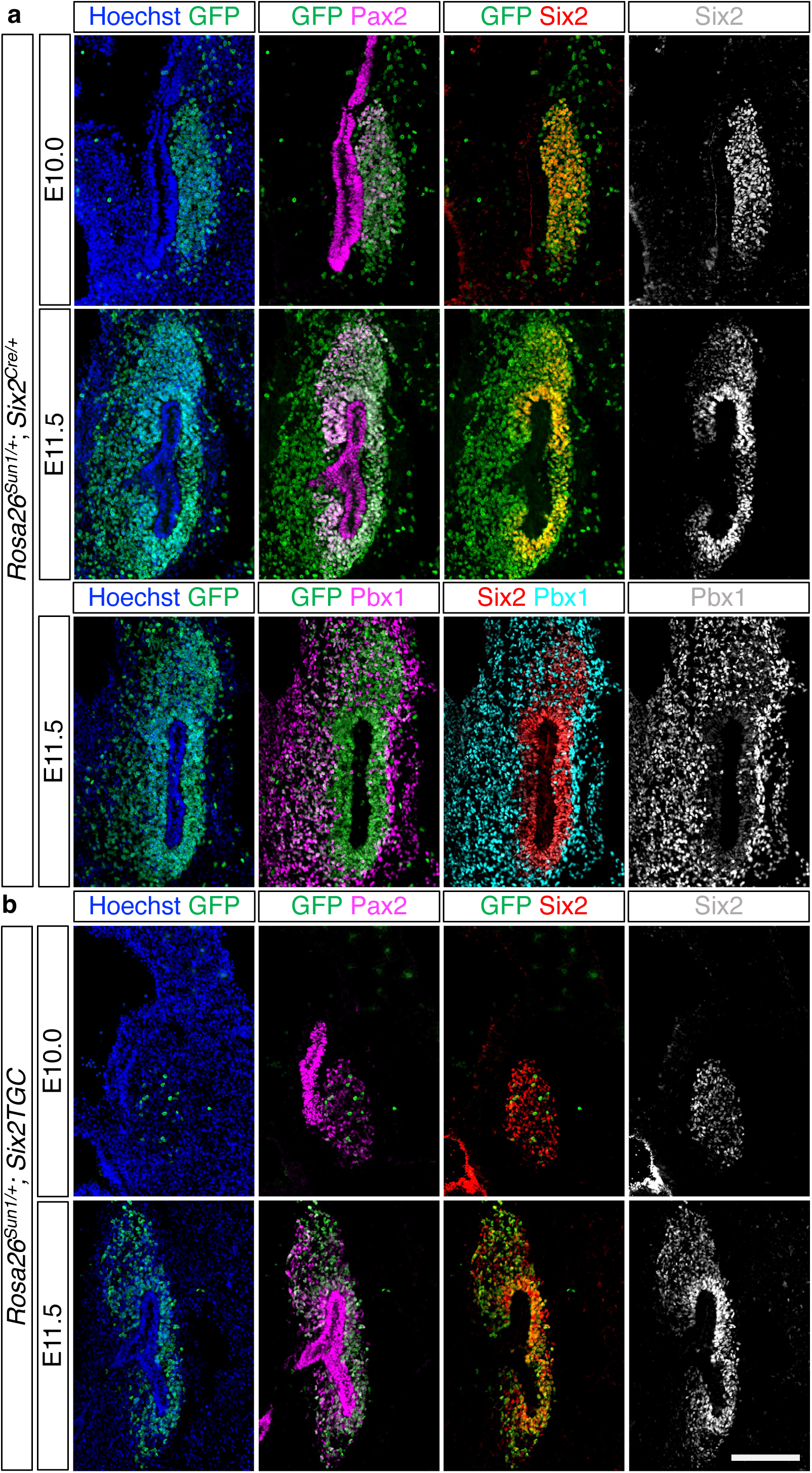
Early *Six2*-expressing cells contribute to nephron and interstitial lineages. The *Rosa26-Sun1* reporter is activated by *Six2Cre* (**a**) or *Six2TGC* (**b**). Pax2 labels nephron and collecting duct lineages, whereas strong Pbx1 marks interstitial cells. (**a**) At E10.0, *Six2Cre* labels all Six2+ Pax2+ mesenchymal nephron progenitor cells adjacent to the nephric duct, which shows high *Pax2* expression. At E11.5, a population of GFP+ Six2- Pax2- cells is observed around the stalk of the branched ureteric epithelium. Adjacent sections show high *Pbx1* expression in these cells. (**b**) In *Six2TGC* kidneys, most Six2+ cells are negative for GFP at E10.0, whereas a majority become positive for GFP by E11.5. Notably, when the reporter is activated by *Six2TGC*, GFP+ Six2- Pax2- interstitial cells are rarely detected. (**a, b**) Scale bar, 100 µm.

By E11.5, however, we observed a distinct population of GFP+ cells that are negative for Six2 or Pax2. These GFP+ Six2- Pax2- cells were largely located on the ventral side of the branched ureteric epithelium (Fig. 4a).

Analysis of adjacent tissue sections revealed that these cells showed high levels of Pbx1, a marker of interstitial lineage, indicating that they are interstitial cells derived from early *Six2*-expressing progenitors.

We performed parallel lineage tracing experiments using the BAC transgenic *Six2TGC* allele. In *Six2TGC* kidneys, most Six2+ cells were not labeled by GFP at E10. By E11.5, the majority of Six2+ cells were GFP+; however, GFP+ Six2- Pax2- cells were rarely observed on the ventral side of the branched ureteric epithelium (Fig. 4b). This contrasts with *Six2Cre* lineage tracing and suggests that delayed recombination in *Six2TGC* kidneys fails to capture interstitial cells derived from *Six2*-expressing mNPs during early kidney development.

Together, these data indicate that *Six2*-expressing cells in the mouse embryonic kidney at E10 have the potential to give rise to both nephron progenitors and interstitial cells. Because genetic labeling of interstitial cells is detected with *Six2Cre* but not with *Six2TGC*, and because *Six2TGC* expression initiates approximately one day later than *Six2Cre*, these findings suggest that Six2+ mNPs largely lose the capacity to generate interstitial cells by E11.5. This finding provides developmental context for the contribution of early mNPs to the renal interstitium.

Although *Six2TGC* has been widely used to trace nephron progenitors, its potential contribution to interstitial lineages has not been systematically examined. Despite its low recombination efficiency at early stages, *Six2TGC* targeted a small number of Six2+ cells at E10 (Fig. 4b). If these early-labeled cells retain interstitial potential, *Six2TGC* lineage tracing would be expected to mark at least a subset of interstitial cells. Consistent with this prediction, activation of the *Rosa26-Sun1* reporter by *Six2TGC* revealed GFP-positive cells with strong Pbx1 signal and no detectable Cdh1 signal in both cortical and medullary regions, indicating interstitial identity (Fig. 5). Together, these findings demonstrate that *Six2TGC* labels a subset of interstitial cells, consistent with early *Six2*-expressing cells retaining interstitial potential.

**Fig. 5:**
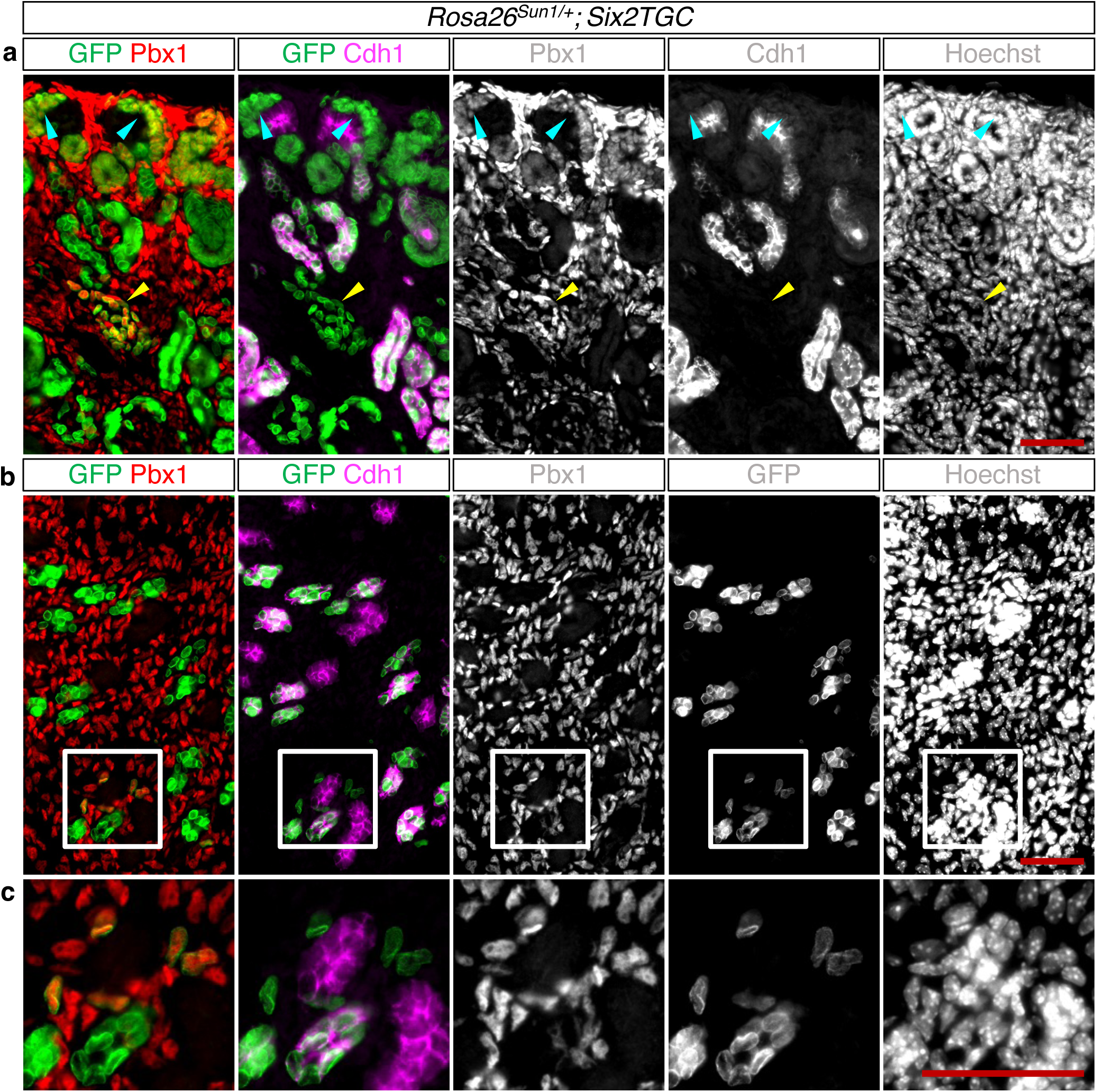
***Six2TGC* targets a subset of interstitial cells** The *Rosa26-Sun1* reporter is activated by *Six2TGC*. Cdh1 labels epithelial cells, and Pbx1 labels non-epithelial cells, including interstitial cells and mesenchymal nephron progenitors. (**a**) In the cortex, cyan arrowheads indicate mesenchymal nephron progenitors, whereas a yellow arrowhead marks a cluster of Pbx1+ Cdh1-interstitial cells labeled by *Six2TGC*. (**b**) In the medulla, a subset of interstitial cells is also labeled by *Six2TGC*. White boxes indicate regions shown at higher magnification in (**c**). (**a-c**) Stage: P0; Scale bar, 100 µm.

#### Early mNPs show a brief trajectory to an interstitial fate

Building on our lineage tracing analyses with *Six2Cre* and *Six2TGC*, which indicate that a subset of early *Six2*-expressing cells contributes to the interstitial compartment, we next sought an independent transcriptomic strategy to test the directionality of this putative transition. To this end, we reanalyzed previously published scRNA-seq data (GSE178263)^26^ from mouse embryonic kidneys and applied RNA velocity analysis^31^ to infer dynamic state changes (Supplementary Fig. 4).

scRNA-seq at E15.5 showed complete segregation of nephron and interstitial lineages: *Six2* was confined to the mNP cluster, whereas *Foxd1* and *Tbx18* marked discrete subsets of interstitium, indicating lineage separation by mid-gestation (Fig. 6a). In contrast, scRNA-seq at E11.5 showed two major clusters that were closely associated, with *Six2* labeling early mesenchymal nephron progenitors and *Foxd1*/*Tbx18* labeling a neighboring interstitial population (Fig. 6b). Notably, the Six2+ and Foxd1+/Tbx18+ clusters were not only positioned in close proximity but also displayed a visible connection in the UMAP embedding. This spatial relationship suggested a developmental link between the two compartments at early stages.

To examine dynamics between these main populations, we performed RNA velocity on the E11.5 dataset. Both grid and stream visualizations revealed coherent vector fields originating within the *Six2*-expressing cluster and projecting toward the *Foxd1*/*Tbx18*-expressing interstitial cluster (Fig. 6c), consistent with a directional transition from mNPs to interstitial identity during a narrow developmental window.

We validated this inference *in vivo* using hybridization chain reaction (HCR)^32^ at E11.5 mouse kidney, in which the *Rosa26*-*Sun1* reporter is activated by *Six2Cre*. GFP-expressing cells were detected within mesenchyme on the ventral side of the branching ureteric epithelium and showed clear co-expression of *Tbx18*, consistent with an interstitial identity (Fig. 6d). The presence of these GFP+ Tbx18+ cells within the interstitial compartment supports the conclusion that a subset of early Six2+ mNPs develops into the Tbx18+ interstitium.

It has been reported that *Tbx18Cre* labels not only ureteric interstitial cells but also cortical stromal progenitors, which are marked by the canonical cortical stroma marker Foxd1, particularly on the ventral side of the kidney adjacent to the ureter^10^. In contrast, our analyses show that *Six2Cre* labels cortical stromal progenitors on both the dorsal and ventral sides with comparable efficiency (Supplementary Fig. 3). This pattern indicates that early Six2+ mNPs contribute to a subset of Foxd1+ cortical stromal cells in addition to Tbx18+ ureteric interstitial cells.

Together, these data indicate that Six2+ mNPs display transient dual potential at E11.5, which resolves into fully segregated nephron and interstitial lineages by E15.5.

## Discussion

In this study, we introduce a knock-in *Six2Cre* allele that faithfully recapitulates endogenous *Six2* expression, maintains nephron endowment, and achieves near-complete, non-mosaic recombination. This new line addresses the major limitations of the widely used BAC transgenic *Six2TGC* allele^3, 7^, which exhibits mosaicism^11, 14^ and reduced nephron number^7, 13, 18, 19^. Using side-by-side lineage tracing with *Six2Cre* and *Six2TGC*, together with RNA velocity analysis of a previously published E11.5 mouse embryonic kidney scRNA-seq dataset, we identify a short developmental interval around E11 during which a subset of Six2+ mNPs contributes to the renal interstitium, including Tbx18+ cells on the ventral side of the branching ureteric epithelium. Notably, mNPs labeled by *Six2Cre* at E10 contribute to interstitial cells more robustly than those labeled by *Six2TGC* at E11.5, indicating that this interstitial contribution is transient and stage-restricted, rather than a persistent feature of Six2+ mNPs.

Our combined lineage tracing and transcriptomic analyses converge on an early developmental trajectory in which Six2+ mNPs give rise to interstitial cells. At E11.5, nephron and interstitial clusters lie in close proximity with visible continuity in the UMAP (Fig. 6b), and RNA velocity vectors extend from the Six2+ cluster toward the Foxd1+/Tbx18+ interstitial population (Fig. 6c), supporting a directional transition during this narrow window. In embryos, in which the *Ros26-Sun1* reporter is activated by *Six2Cre*, we found GFP+ Tbx18+ cells within interstitial cells adjacent to the stalk of the ureteric epithelium (Fig. 6d), providing spatial validation of this early lineage relationship. By E15.5, nephron and interstitial clusters are fully segregated (Fig. 6a), indicating that the early plasticity observed at E11.5 resolves into stable, committed nephron and interstitial lineages.

**Fig. 6:**
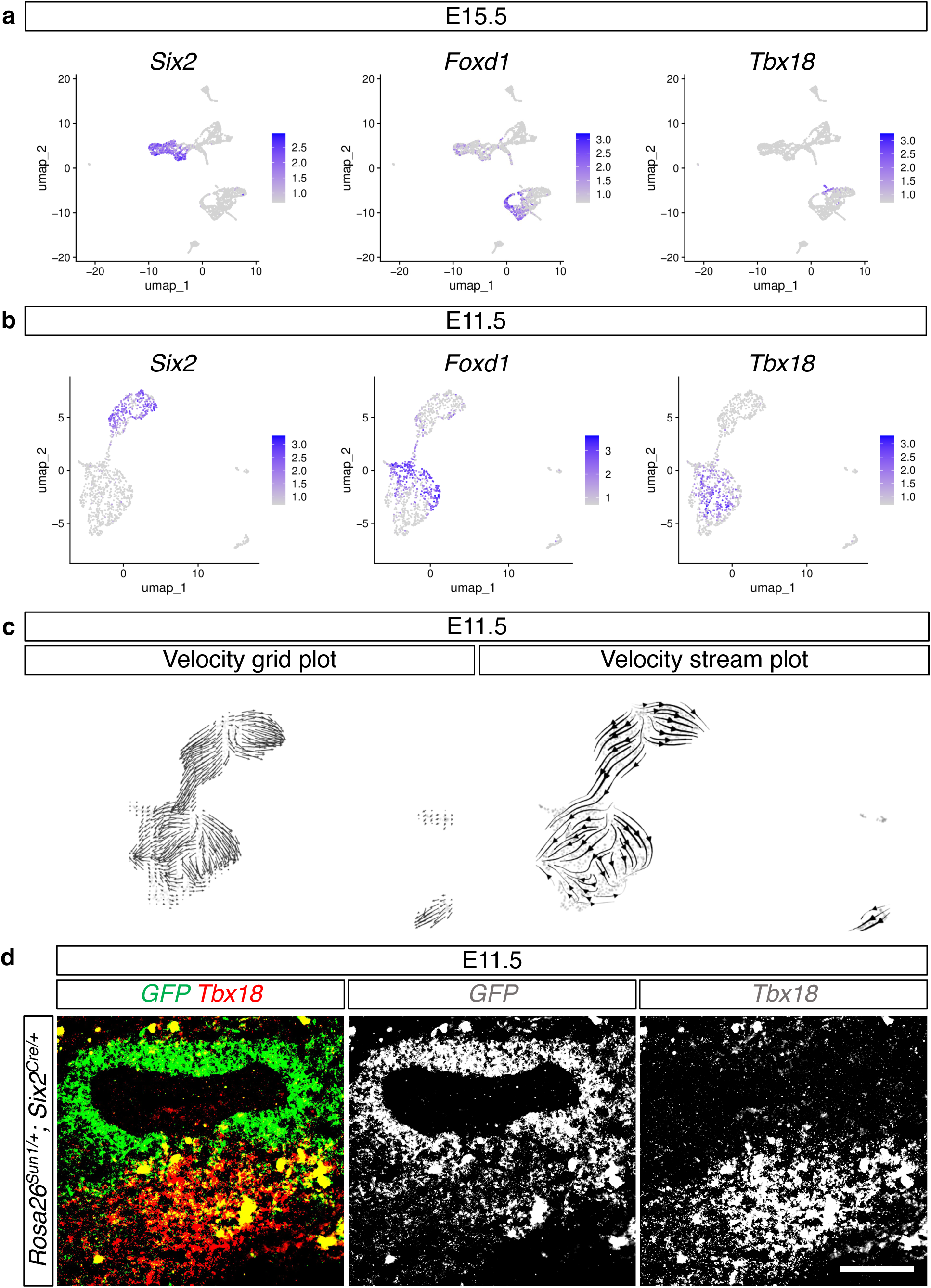
Early nephron progenitors exhibit a transient trajectory toward an interstitial fate. **(a)** Feature plots of *Six2*, *Foxd1*, and *Tbx18* expression on E15.5 scRNA-seq UMAPs show complete segregation of nephron progenitor and interstitial lineages. (**b**) At E11.5, *Six2* labels mesenchymal nephron progenitors, while *Foxd1* and *Tbx18* label adjacent interstitial populations positioned in close proximity in UMAP space. (**c**) RNA velocity analysis of E11.5 scRNA-seq data reveals vector flows extending from the Six2+ nephron progenitor cluster toward the Foxd1+/Tbx18+ interstitial cluster. (**d**) Hybridization chain reaction detects *GFP* expression from *Six2Cre*-activated Rosa26-Sun1 reporter together with endogenous *Tbx18* expression in cells located on the ventral side of the branched ureteric epithelium. Stage: E11.5; Scale bar, 100 µm.

These observations align with and extend prior mammalian lineage tracing studies that challenge strict epithelial-stromal boundaries. *Foxd1Cre*, expressed earlier and more broadly (including in somites), marks interstitial cells but also small populations of mNPs (Fig. 6a), a pattern often attributed to “driver leak” but that may instead reflect true biological adjacency or limited early lineage flux. *Six2Cre* is specifically expressed in mNPs yet still detects a Six2-derived contribution to Tbx18+ interstitium, complementing *Foxd1Cre*’s stromal-first perspective and reinforcing the concept of transient mesenchymal plasticity at compartment interfaces.

The transient interstitial contribution of early Six2+ mNPs is further supported by genetic studies demonstrating that nephron progenitor identity must be actively maintained. Loss of Pax2, a core nephron lineage determinant, has been shown to induce transdifferentiation of mNPs into interstitial cell types^33^, revealing an inherent susceptibility of these cells to adopt a stromal fate when nephron programs are compromised. These findings indicate that the nephron-interstitium boundary is enforced by transcriptional regulation rather than irrevocable lineage commitment, and they provide a mechanistic precedent for the early, stage-restricted plasticity uncovered by our study.

Tbx18 promotes ureteric interstitium specification by repressing metanephric mesenchymal programs and delineating the mesenchymal domain around the distal ureteric bud^34^. Earlier work places Tbx18 within a urogenital mesenchyme patterning region that biases ureter versus cortical stromal identity^10^. Interpretation of Tbx18 lineage contributions depends on the timing of Cre activation. When an inducible *Tbx18Cre* is activated before E11.5, it labels Foxd1+ stromal cells due to a transient Foxd1/Tbx18 double-positive population; in contrast, activation at E12.5 does not label Foxd1+ stromal cells^35^. Together, these observations position our *Tbx18*-expressing cells arising from the Six2 lineage within this early patterning zone, where local cues can transiently bias the fate of mNPs toward Foxd1/Tbx18 interstitial lineages.

Viewed in a broader, comparative context, cross-species evidence further supports a flexible view of mesodermal contributions to kidney lineages. In zebrafish, somites derived from paraxial mesoderm can contribute to nephrons^36^, challenging the long-standing assumption that kidneys derive exclusively from intermediate mesoderm^37, 38^. Although species differences and organ architecture prevent direct equivalence with the mouse, the underlying concept of permissive interfaces between mesodermal territories that allow limited, context-dependent lineage flux is consistent with the early transition from *Six2*-expressing mNPs to interstitial cells observed here.

In sum, we present a knock-in *Six2Cre* allele that enables precise, non-mosaic targeting of nephron progenitors and reveals that early Six2+ mNPs exhibit transient dual potential, contributing to an interstitial lineage during a narrow developmental window before separating into fully segregated nephron and stromal lineages. When placed alongside lineage-tracing studies of renal interstitial cells^8, 10, 34, 35, 39^ and zebrafish evidence for somite contributions to nephrons^36^, these findings support a plastic, context-dependent view of kidney lineage specification. By clarifying early inter-compartment relationships and providing a robust genetic entry point into nephron progenitors, this work establishes a framework for dissecting how timing, niche signals, and transcriptional state coordinate nephron and stromal development, and it suggests avenues to leverage early plasticity in regenerative strategies.

## Methods

### Generation of *Six2* knock-in Cre

CRISPR/Cas9 genome editing was performed by the Transgenic Animal and Genome Editing Core at Cincinnati Children’s Hospital Medical Center. A DNA fragment encoding T2A-NLS-GFP-Cre was inserted immediately downstream of the Six2 stop codon (Supplementary Fig. 1). A single guide RNA (sgRNA) was selected based on optimal on-target and off-target scores predicted by CRISPOR^40^. The donor plasmid was generated by cloning synthesized DNA fragments (Gene Universal) into the pBluescript vector, containing *T2A*1J*NLS*1J*GFP*1J*Cre* flanked by 2.0 kb 5′ and 2.1 kb 3′ homology arms. The donor plasmid was amplified and purified using the EndoFree Plasmid kit (Qiagen). Fertilized C57BL/6J zygotes were injected with Cas9 protein (IDT, 1081061; 40 ng/µl), sgRNA (30 ng/µl), and donor plasmid (5 ng/µl), and transferred into pseudo-pregnant CD-1 females. Founder mice were genotyped by long-range PCR and Sanger sequencing. The sequences of oligonucleotides P1-P7 are shown in Supplementary Table 1.

#### Mice

*Hnf4a flox*^41^, *Rosa26-Sun1* (JAX:021039)^27^, and *Six2TGC* (JAX:009606)^3, 7^ have been previously reported. All animal studies were conducted in accordance with animal care guidelines, and the protocols were approved by Institutional Animal Care and Use Committees at Northwestern University and Cincinnati Children’s Hospital and Medical Center, in accordance with the relevant NIH Guide for the Care and Use of Laboratory Animals.

Animals were housed in a controlled environment under a 12-h light/12-h dark cycle, with ad libitum access to water and a standard chow diet. Mice of both sexes were included in the study, and no sex-dependent phenotypes were observed.

#### Tissue processing

Mouse embryos were examined for the *Six2*-driven *GFP* expression pattern using a fluorescence microscope (Leica 2, MDG41; Fluorescent source Prior Lumen 200, L200US) equipped with Leica 2 microsystem CMS (D35578, type DFC310FX) camera using Leica 2 Application Suite X 3.7.0.20979. Whole embryos or dissected kidneys were fixed in 4% paraformaldehyde (Electron Microscopy Science) in phosphate buffered saline (PBS) for 20 minutes (for immunofluorescence staining) or overnight (for hybridization chain reaction). Fixed tissues were incubated overnight in 10% sucrose in PBS, embedded in an optimal cutting temperature compound (Tissue-Tek O.C.T; Sakura), and quickly frozen on a dry ice-ethanol bath. Frozen blocks were stored at −80°C until sectioning (8-10 µm) using a Epredia Cryostar NX50 cryostat.

#### Fluorescence-activated cell sorting

E18.5 mouse embryonic kidneys carrying either *Six2TGC* or *Six2Cre* were minced by razor blades, and digested in TrypLE Select Enzyme (Gibco) by periodic pipetting at 37°C for 20 minutes. Dissociated cells were suspended in PBS containing 10% fetal bovine serum (FBS)(Gibco) and loaded onto a 40 µm nylon cell strainer (BD Falcon). Filtrates are centrifuged (Beckman Coulter, Allegra X-30 R; 300x g, 5 minutes at 4°C), and resuspended cell pellets were incubated in red blood cell lysis buffer (Hybrid-Max, Sigma life science) for 2 minutes, continued by a sequential washing in 1X PBS containing 10% FBS and centrifugation steps (Eppendorf centrifuge 5417R; 300 xg, 5 minutes at 4°C). Pellets are then resuspended in PBS containing 1% FBS and 10 mM EDTA (Thermo Fisher Scientific). Cells were kept on ice throughout the procedure except for the initial digestion. GFP-positive cells were isolated using a BD FACSAria II, spun down, and stored at −80°C.

#### Bulk RNA-sequencing

RNA extraction was performed using Single Cell RNA Purification Kit (Norgen Biotech). NEBNext Poly(A) mRNA Magnetic Isolation Module (E7490, New England Biolabs) was used to purify mRNAs from total RNA. cDNA synthesis was performed using NEBNext RNA First (E7525L) and Second Strand (E6111L) Synthesis Modules (New England Biolabs). Sequencing libraries were prepared using the ThruPLEX DNA-seq 12S Kit (R400428, Takara Bio) and sequenced on either an Illumina NovaSeq 6000 or an Element AVITI platform.

Sequencing reads were aligned to the mouse reference genome (mm10) using HISAT2^42^, and differential gene expression analysis was conducted using Cuffdiff (v2.1.1)^43^. Genes exhibiting at least a 1.5-fold change in expression with *P* < 0.05 were considered differentially expressed.

#### Immunofluorescent staining

Cryosections were washed with PBS and PBST (PBS containing 0.1% Triton X-100), and incubated with 5% heat-inactivated sheep serum (Equitech-Bio, Inc) in PBST for 1 hour. Sections were incubated with the primary antibody mix overnight at 4°C. After washing steps (PBST), they were incubated with secondary antibodies (1:500) and Hoechst DNA stain (1:2000; Invitrogen) for 1 hour and 15 minutes, respectively, at room temperature. Mounting media (P36930, Prolong Gold antifade reagent, ThermoFisher Scientific) was applied to each section after additional washing steps in PBST and a brief wash in deionized water, followed by sealing with a coverslip (UWR coverslip, 22×40 mm, 1.5). Details of all primary and secondary antibodies used in this study are provided in Supplementary Table 2. Imaging was performed using a Nikon Ti2 Eclipse inverted wide-field microscope equipped with a Spectra light engine (Lumencor Spectra III light source) and a camera (Orca Fusion). Image J (v1.53) was used for image processing.

#### Hybridization chain reaction (HCR)

HCR was performed as described previously with some modifications^32^. Cryosections were washed in 1X PBS and exposed to a combination of the photo (Gooseneck LED desk lamp; output: 5V-2A) and H_2_O_2_ (C762T82, H1070-100ML-CS6, Spectrum Chemicals)-based chemical auto-fluorescent bleaching solution (Supplementary Table 3) at 4°C for 1 hour. Following washing steps (1X PBS, 3 x 10 minutes, PBS/Tween 20, 3 x 5 minutes, 5X Saline Sodium Citrate /Tween 20, 3 x 5 minutes, and a quick rinse with 1X PBS at room temperature), sections were incubated in probe hybridization buffer for 30 minutes at 37°C, followed by overnight incubation with ssDNA probe sets for GFP and Tbx18 at 37°C. The sections were incubated in the probe wash buffer at 37°C for 30 minutes. After two additional rounds of washes with probe wash buffer (2 × 15 minutes; 37°C) and 5X SSC-T (3 × 5 minutes; room temperature), the sections were incubated in amplification buffer for 30 minutes at room temperature. The buffer was replaced with amplification buffer containing amplifier DNA hairpins for GFP (1:50 dilution) and Tbx18 (1:25 dilution). Then, the slides were washed sequentially with 1X PBS (5 minutes) and 5X SSC-T (1 x 5 minutes, 2 x 30 minutes) in a dark drawer. Hoechst DNA stain (10 minutes; 1:2000; Invitrogen) was used to mark cell nuclei. After mounting, slides were kept at room temperature and protected from light before imaging. Detailed information regarding the target genes, probes, and hairpins is provided in Supplementary Table 4.

#### Single-cell RNA-seq and RNA velocity analysis

scRNA-seq data from mouse embryonic kidneys at E11.5 and E15.5 (GSE178263)^26^ was obtained from Gene Expression Omnibus. FASTQ files were processed using the Cell Ranger pipeline (v8.0.1, 10x Genomics) with the mm10 reference genome. All downstream analyses were performed in R (v4.4.0) using Seurat (v5.3.0)^44^. Quality filtering was performed based on UMI counts, detected genes, and mitochondrial content (18,000 – 75,000 UMIs; 5,000 – 9,000 genes; < 5.5% mitochondrial). Cells outside these ranges were excluded from further analysis. Data were normalized using SCTransform (v2) with the glmGamPoi method. Cell clustering was performed using the Louvain algorithm of the FindClusters function with a resolution of 0.4. Uniform Manifold Approximation and Projection (UMAP) dimensionality reduction was computed using the first 30 principal components. Cluster marker genes were found using the FindAllMarker function with parameters logfc.threshold > 0.25 and min.pct > 0.1. Differential genes were determined using the FindMarkers function with parameters logfc.threshold > 0.5 and min.pct > 0.5. For RNA velocity analysis, spliced and unspliced transcript counts were generated from the BAM file using velocyto^31^. RNA velocity estimation was performed using scVelo^45^ in stochastic mode, and velocity vectors were projected onto the UMAP embedding using both stream and grid plots.

## Data availability

The bulk RNA-seq data from this study were deposited in the Gene Expression Omnibus (GEO) under accession numbers GSE314255.

## Acknowledgements

This work was supported by National Institutes of Health, National Institute of Diabetes and Digestive and Kidney Diseases grants DK125577, DK131052, DK127634, DK120847, and DK120842.

## Author Contributions

Conceptualization: A.H., E.C., J.S.P.

Methodology: A.H., Y.C.H., E.C.

Validation: A.H.

Formal analysis: M.S., C.A., H.W.L., J.S.P.

Investigation: A.H., F.N., Z.D.G., E.C., J.S.P.

Resources: J.S.P.

Data curation: A.H., J.S.P.

Writing – original draft: A.H.

Writing – review and editing: E.C., J.S.P.

Visualization: A.H.

Supervision: J.S.P.

Project administration: J.S.P.

Funding acquisition: J.S.P.

## Supplementary information

Supplementary Figure 1. Generation of the *Six2* knock-in Cre allele

Supplementary Figure 2. Bulk RNA-seq analysis of FACS-isolated mNPs

Supplementary Figure 3. Lineage tracing with Six2Cre in the E15.5 mouse kidney

Supplementary Figure 4. Single-cell transcriptomic landscapes of mouse embryonic kidneys (GSE178263)

Supplementary Table 1. Oligonucleotides used to genotype the *Six2* knock-in Cre allele

Supplementary Table 2. Primary and secondary antibodies used in this study

Supplementary Table 3. HCR RNA FISH auto-fluorescence bleaching solution components

Supplementary Table 4. HCR RNA FISH probes and amplifiers

