## Supplementary Figures 1-4 for "A knock-in *Six2Cre* line reveals transient interstitial potential in nephron progenitors"

**a** *Six2* genomic locus

**b** Targeted locus

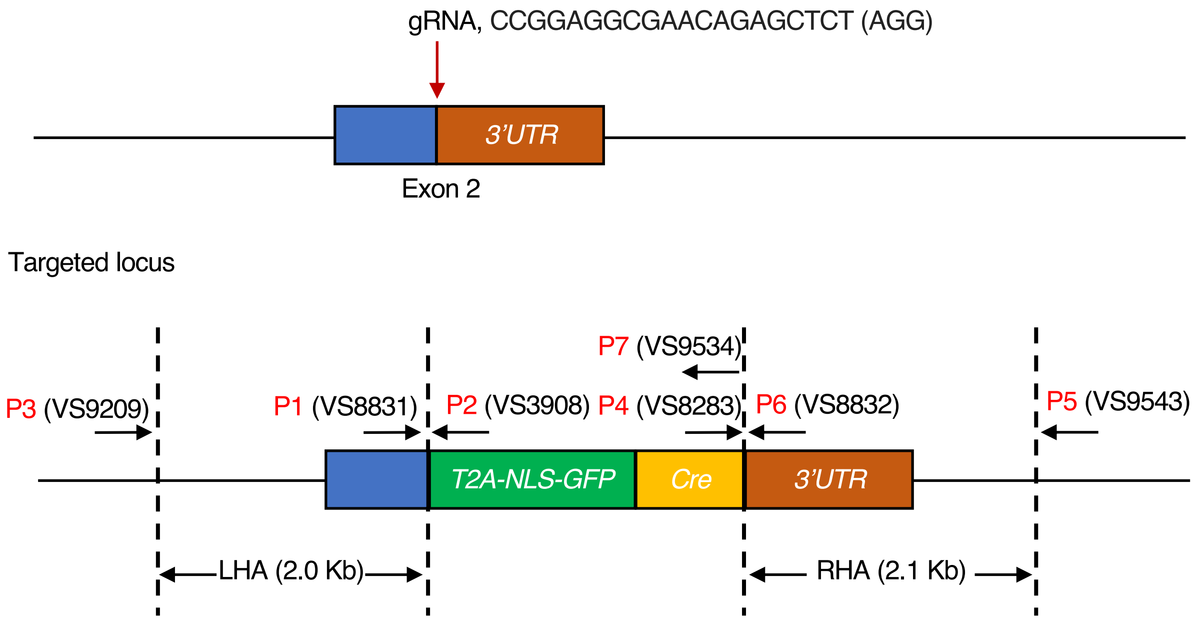


**Supplementary Figure 1. Generation of the *Six2* knock-in Cre allele**

(**a**) Schematic of the mouse *Six2* genomic locus, showing the gRNA target site adjacent to the Six2 stop codon. The adjacent AGG sequence denotes the protospacer-adjacent motif (PAM).

(**b**) Diagram of the CRISPR/Cas9-engineered knock-in allele. A T2A–NLS–GFP–Cre cassette was inserted immediately upstream of the Six2 stop codon. Primer sequences (P1–P7) used to confirm correct targeting are listed in Supplementary Table 1.

Abbreviations: gRNA, guide RNA; NLS, nuclear localization signal; UTR, untranslated region; LHA, left homology arm; RHA, right homology arm.

**a**

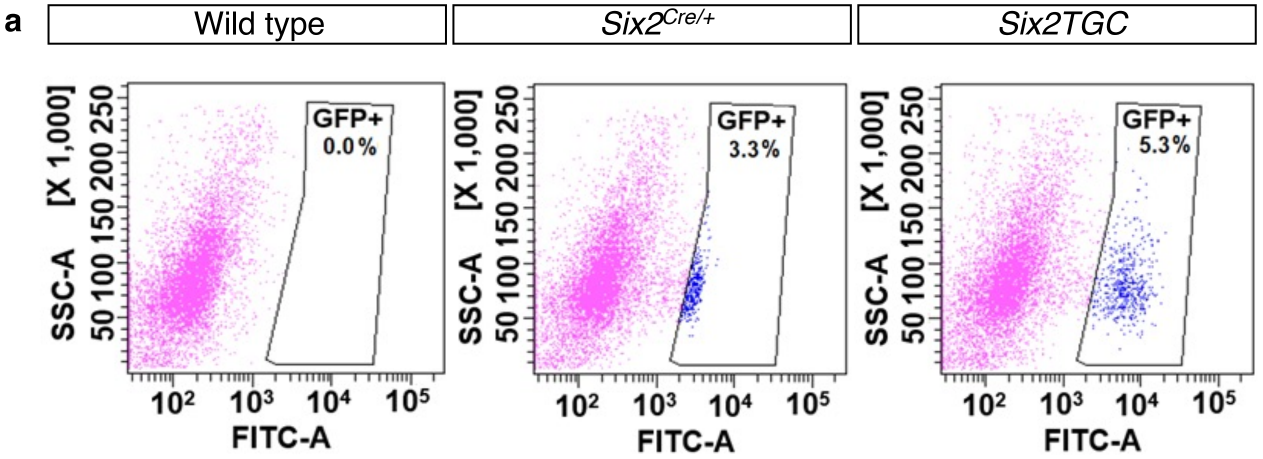


**b**

| Gene | log2(FC) | P-value | Avg_Six2TGC | Avg_Six2Cre | Status | Six2TGC_1 | Six2TGC_2 | Six2Cre_1 | Six2Cre_2 |
|---|---|---|---|---|---|---|---|---|---|
| **Transcriptional regulators expressed in mesenchymal nephron progenitors** | | | | | | | | | |
| *Six2* | -0.54 | 0.0024 | 782.44 | 539.00 | Unchanged | 748.69 | 816.20 | 544.98 | 533.03 |
| *Six3* | - | 0.0001 | 106.15 | 0.00 | Down | 106.69 | 105.61 | 0.00 | 0.00 |
| *Eya1* | 0.29 | 0.0604 | 73.02 | 89.41 | Unchanged | 69.13 | 76.91 | 91.09 | 87.73 |
| *Osr1* | -0.01 | 0.9497 | 60.98 | 60.53 | Unchanged | 52.61 | 69.34 | 58.73 | 62.33 |
| *Pax2* | 0.50 | 0.0017 | 158.00 | 223.33 | Unchanged | 148.70 | 167.30 | 231.08 | 215.58 |
| *Wt1* | 0.36 | 0.0187 | 119.19 | 153.24 | Unchanged | 114.46 | 123.91 | 154.48 | 152.00 |
| *Sall1* | -0.17 | 0.2853 | 184.55 | 163.58 | Unchanged | 179.35 | 189.74 | 161.58 | 165.58 |
| *Cited1* | 3.13 | 0.0001 | 101.75 | 892.20 | Up | 108.92 | 94.58 | 906.86 | 877.55 |
| **Genes activated during mesenchymal-to-epithelial transition** | | | | | | | | | |
| *Wnt4* | -3.48 | 0.0001 | 114.92 | 10.28 | Down | 110.07 | 119.77 | 9.36 | 11.20 |
| *Jag1* | -1.57 | 0.0001 | 12.46 | 4.18 | Down | 11.69 | 13.22 | 3.80 | 4.57 |
| *Notch1* | -0.91 | 0.0001 | 23.07 | 12.25 | Down | 25.25 | 20.88 | 12.90 | 11.60 |
| *Lhx1* | -2.75 | 0.0001 | 7.33 | 1.09 | Down | 8.00 | 6.66 | 1.01 | 1.18 |
| *Fgf8* | 2.07 | 0.0001 | 9.64 | 40.36 | Up | 10.05 | 9.22 | 39.44 | 41.29 |

**c**

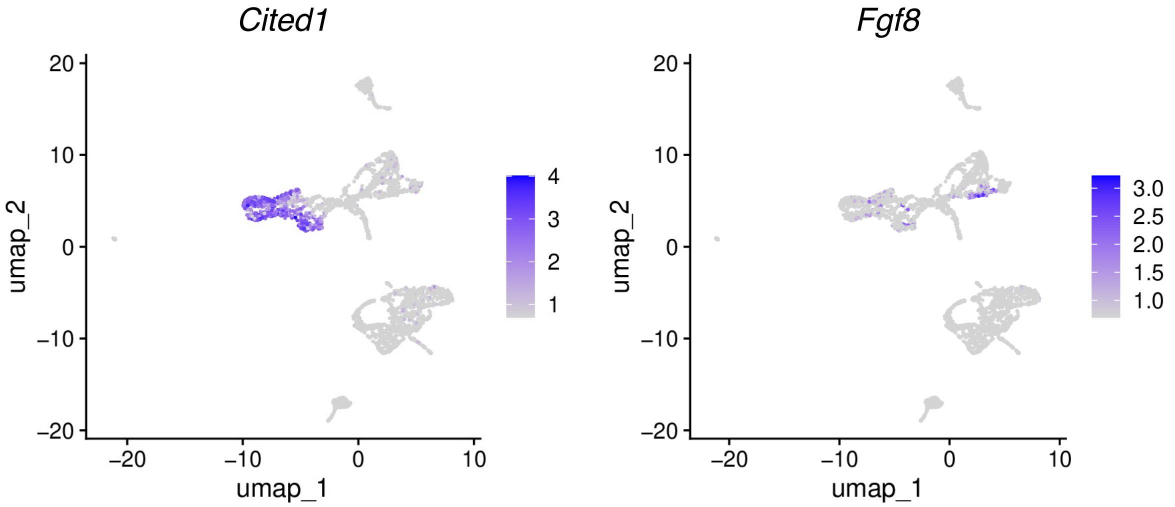


**Supplementary Figure 2. Bulk RNA-seq analysis of FACS-isolated mNPs**

(**a**) Gating strategy used to isolate GFP+ cells from E18.5 mouse embryonic kidneys. The collected GFP+ population (shown in blue) was used for bulk RNA-seq.

(**b**) Expression of selected genes in mNPs expressing *Six2TGC* or *Six2Cre*. FC indicates the fold change (*Six2Cre*/*Six2TGC*).

(**c**) Single-cell RNA-seq data from E15.5 mouse embryonic kidneys (GSE178263) showing that *Fgf8* expression is detectable within Cited1+ mNPs.

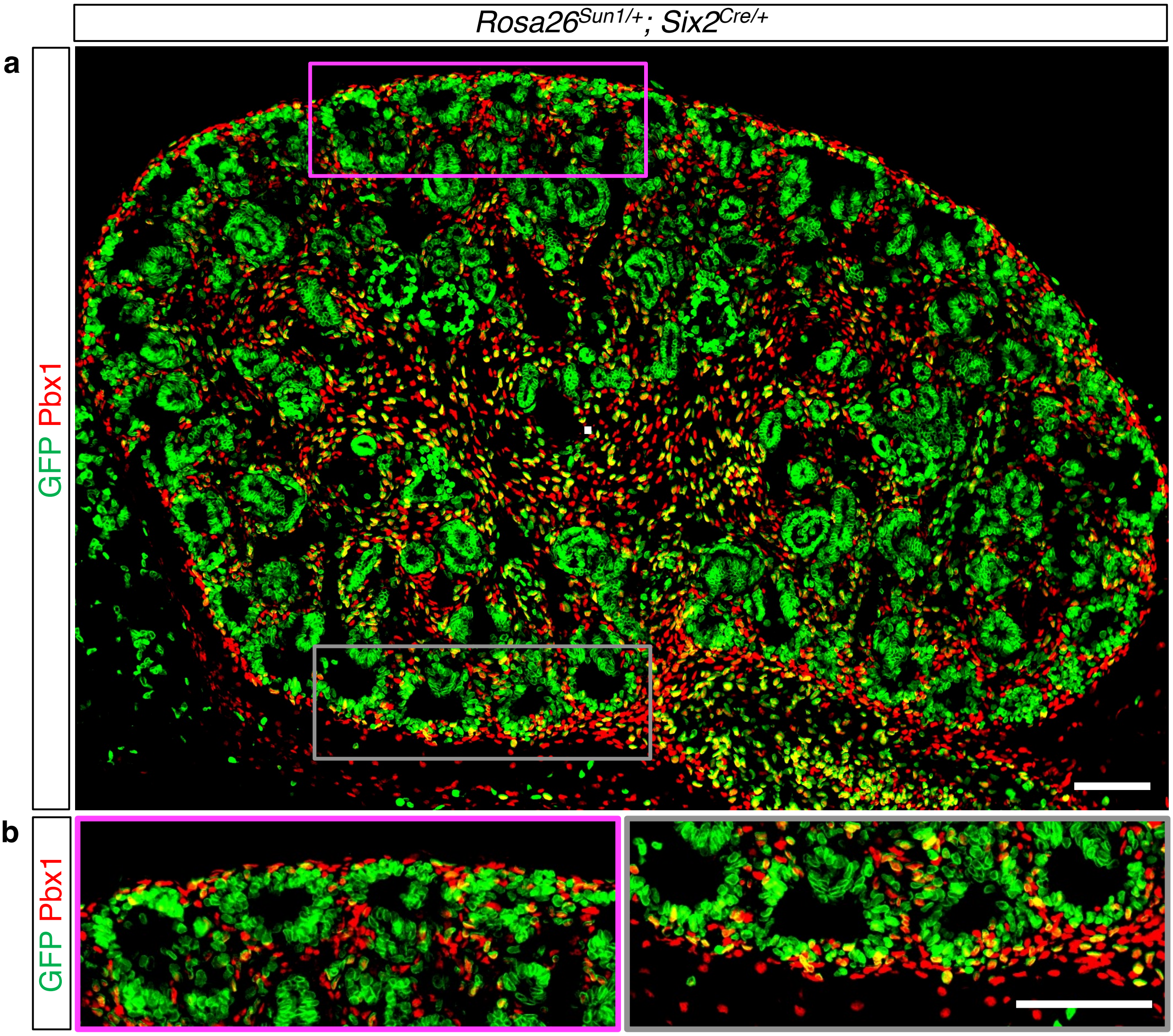


**Supplementary Figure 3. Lineage tracing with *Six2Cre* in the E15.5 mouse kidney**
(**a**) Immunofluorescence analysis showing Pbx1+ interstitial cells labeled by *Six2Cre*-mediated activation of the *Rosa26-Sun1* reporter. GFP+ Pbx1+ cells are present throughout the E15.5 mouse embryonic kidney.
(**b**) Higher-magnification views of the dorsal cortex (magenta box) and ventral cortex (grey box).(**a**-**b**) Stage, E15.5; Scale bar, 100 $\mu$m.

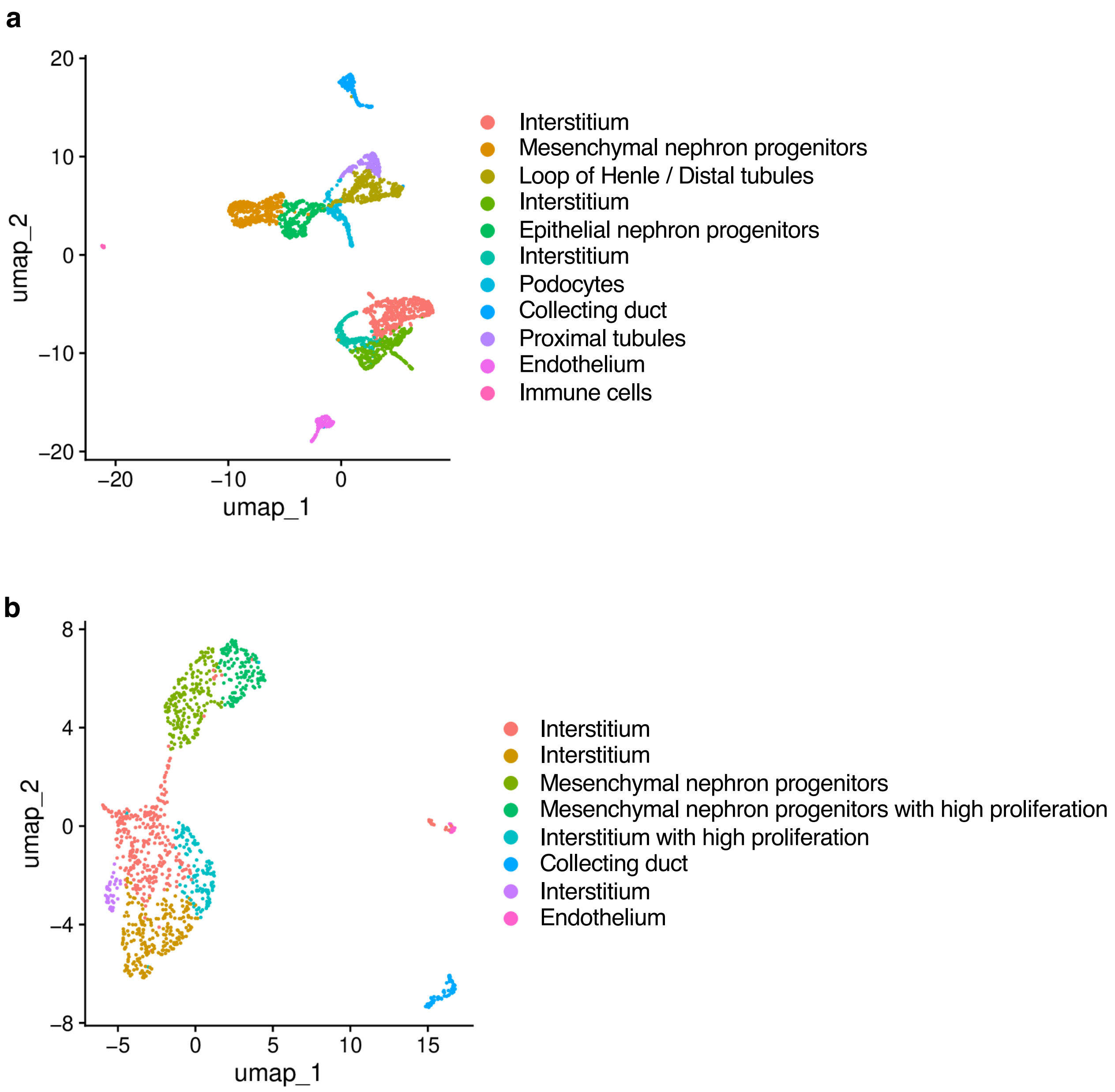


**Supplementary Figure 4. Single-cell transcriptomic landscapes of mouse embryonic kidneys (GSE178263)**
(**a**) UMAP projection of single-cell RNA-seq data from E15.5 mouse kidneys, showing clear segregation between nephron and interstitial lineages.
(**b**) UMAP projection of single-cell RNA-seq data from E11.5 mouse kidneys, in which nephron progenitors and interstitial populations are positioned in close proximity.
