## Supplementary Tables 1-4 for "A knock-in *Six2Cre* line reveals transient interstitial potential in nephron progenitors"

**Supplementary Table 1.** Oligonucleotides used to genotype *Six2* knock-in Cre allele

| **Primers** | **Sequence (5’ – 3’)** | **Forward/ Reverse** |
| --- | --- | --- |
| P1 (VS8831) | GACCCACTGCAGCATCACCAC | Forward |
| P2 (VS3908) | TGAACTTGTGGCCGTTTACGTC | Reverse |
| P3 (VS9209) | AGCTCCTGGACAGGACTAAATCGC | Forward |
| P4 (VS8283) | GATGACTCTGGTCAGAGATACCTG | Forward |
| P5 (VS9543) | AGGGACCCCAAGGGCCCAAGGATACG | Reverse |
| P6 (VS8832) | TGGCAGGGCAGTCTTCCAGTAC | Reverse |
| P7 (VS9534) | AAACTAATCGCCATCTTCCAGCAGGC | Reverse |

***Note***: F1 pups were genotyped using both internal and external primers (Supplementary Fig. 1), as the founders can carry both targeted and randomly integrated transgenes. Copy number analysis was used to ensure that the F1 pups have a single copy of the transgene. Starting from F2, pups were genotyped routinely with a 3-primer strategy (primers P1, P2, and P6; Cre: 247 bp; WT: 185 bp).

**Supplementary Table 2.** Primary and secondary antibodies

| **Item** | **Targets** | **Class** | **Host** | **Type** | **Suppliers** | **Catalog #** | **Dilution** |
| --- | --- | --- | --- | --- | --- | --- | --- |
| 1 | GFP | Ig | Guinea pig | Polyclonal | Synaptic Systems | 132 005 | 1:6000 |
| 2 | Pax2 | IgG | Rabbit | Polyclonal | Covance | PRB-276P | 1:200 |
| 3 | SIX2 (clone: 3D7) | IgG1 | Mouse | Monoclonal | Novus Biologicals | H00010736-M01 | 1:500 |
| 4 | Pbx1 | IgG | Rabbit | Polyclonal | Proteintech | 18204-1-AP | 1:500 |
| 5 | Cdh1  (E-cadherin) | IgG | Rat | Monoclonal | Santa Cruz | SC-59778 | 1:500 |
| 6 | Hnf4a | IgG | Rabbit | Polyclonal | Santa Cruz | sc-8987 X | 1:500 |
| 7 | FITC-Lotus Tetragonolobus Lectin (LTL) | - | - | - | Vector labs | FL-1321 | 1:200 |
| 8 | Alexa Fluor® 488 AffiniPure Donkey Anti-Guinea Pig IgG (H+L) | IgG | Guinea | - | Jackson Immuno Research Labs | 706-545-148 | 1:500 |
| 9 | Cy3 Donkey Anti-Rabbit IgG: Rabbit red | IgG | Donkey | - | Jackson Immuno Research Labs | 711-166-152 | 1:500 |
| 10 | Alexa Fluor® 647 Goat Anti-Mouse IgG1 (γ1) | IgG1 (γ1) | Goat | - | Invitrogen | A-21240 | 1:500 |
| 11 | Alexa Fluor® 647 AffiniPure Donkey Anti-Rat IgG (H+L) | IgG | Donkey | - | Jackson Immuno Research Labs | 712-605-153 | 1:500 |

**Supplementary Table 3**: HCR RNA FISH auto-fluorescence Bleaching solution

| **Reagent** | **Final concentration** | **Amount for 3 ml** |
| --- | --- | --- |
| 10X PBS | 1x | 300 µl |
| NaOH 10 N | 24mM | 7 µl |
| 30% H2O2 | 4.5% | 450 µl |
| Water | - | 2243 µl |

**Supplementary Table 4.** HCR RNA FISH probes and amplifiers

| **A. HCR probes** | | | | | | | | |
| --- | --- | --- | --- | --- | --- | --- | --- | --- |
| **Organism** | **Target RNA** | **Accession number/sequence** | **Designed for** | **Probe set size** | **Conc.** | **The amount used/experiment*** | **Length of RNA (nt)** | **Lot No.** |
| Transgenic | d2eGFP | AUggUgAgCAAgggCgAggAgCUgUUCACCggggUggUgCCCAUCCUggUCgAgCUggACggCgACgUAAACggCCACAAgUUCAgCgUgUCCggCgAgggCgAgggCgAUgCCACCUACggCAAgCUgACCCUgAAgUUCAUCUgCACCACCggCAAgCUgCCCgUgCCCUggCCCACCCUCgUgACCACCCUgACCUACggCgUgCAgUgCUUCAgCCgCUACCCCgACCACAUgAAgCAgCACgACUUCUUCAAgUCCgCCAUgCCCgAAggCUACgUCCAggAgCgCACCAUCUUCUUCAAggACgACggCAACUACAAgACCCgCgCCgAggUgAAgUUCgAgggCgACACCCUggUgAACCgCAUCgAgCUgAAgggCAUCgACUUCAAggAggACggCAACAUCCUggggCACAAgCUggAgUACAACUACAACAgCCACAACgUCUAUAUCAUggCCgACAAgCAgAAgAACggCAUCAAggUgAACUUCAAgAUCCgCCACAACAUCgAggACggCAgCgUgCAgCUCgCCgACCACUACCAgCAgAACACCCCCAUCggCgACggCCCCgUgCUgCUgCCCgACAACCACUACCUgAgCACCCAgUCCgCCCUgAgCAAAgACCCCAACgAgAAgCgCgAUCACAUggUCCUgCUggAgUUCgUgACCgCCgCCgggAUCACUCUCggCAUggACgAgCUgUACAAgUAAAgCggCCgCgACUCUAgAUCAUAAUCAgCCAUACCACAUUU | Amplifier B3 | 12 | 1 µM | 1 µl | 761 nt | RTL327 |
| Mouse | Tbx18 | [NM_023814.4](https://www.ncbi.nlm.nih.gov/nuccore/NM_023814.4) | Amplifier B1 | 20 | 1 µM | 2 µl | 4679 nt | RTE292 |
| **B. HCR amplifiers** | | | | | | | | |
| **HCR Amplifier** | | **Amplifier fluorophore** | **Amplifier scale** | **Conc.** | | **Lot No.** | | |
| B3-h1-546 | | 546-red | 600 pmol | 3 µM | | SO94426 | | |
| B3-h2-546 | | 546-red | 600 pmol | 3 µM | | SO94526 | | |
| B1-h1-647 | | 647-farred | 600 pmol | 3 µM | | SO96326 | | |
| B1-h2-647 | | 647-farred | 600 pmol | 3 µM | | SO96426 | | |
